# NaCl Triggers the Sessile-to-Motile Transition of *Bacillus subtilis*

**DOI:** 10.1101/2022.02.15.480532

**Authors:** Prem Anand Murugan, Manish Kumar Gupta, T. Sabari Sankar, Sivasurender Chandran, Saravanan Matheshwaran

## Abstract

Various chemical cues are known to alter the motile and sessile states of bacteria differentially and, in turn, the formation of biofilms. However, the underlying mechanisms at the cellular and molecular level remain less understood, which severely limits our ability to control biofilms. Here, we systematically studied the effects of NaCl on the dynamics of biofilm formation across various length scales and the associated changes in the regulation of gene expression in an undomesticated natural isolate of *Bacillus subtilis*. Interestingly, NaCl induced significant changes in the architecture of pellicles and yielded systematic increase in lateral expansion rates of biofilms when grown on an agar surface. At the microscopic level, both in the presence and absence of NaCl, bacteria displayed super-diffusive motion at times lesser than a second. However, at larger delay times, we observed an intriguing NaCl-induced transition from sub-diffusion behavior of individual bacterial cells to rapid diffusion behavior. In addition, NaCl reduced the dynamical heterogeneity of the bacterial cells within the biofilm. The reduced heterogeneity and the increased flagellation in a subpopulation of cells in the presence of NaCl corroborates well with the observed higher motility of the cells. Further, the cellular uptake of NaCl resulted in the downregulation of several genes underlying the formation of biofilms, revealing the role of chemical cues like NaCl in controlling the gene regulatory circuit underlying the sessile to motile transition. Our study opens a new avenue to decipher the competitive advantage provided to the subcellular populations by NaCl due to lifestyle switch in *Bacillus subtilis*.

## Introduction

Bacteria are the most abundant and diverse form of life on the planet. In their natural environments, they often live in structurally complex and dynamic multicellular communities called biofilms, where bacteria are embedded in the self-secreted viscoelastic fluids comprised of various macromolecules, including exopolysaccharides (EPS), and proteins that stabilize biofilms (1-4). In contrast to the planktonic state, the multicellular lifestyle provides fitness advantages for the microbes by enhancing their tolerance/resistance to external chemical and mechanical stressors (1-4). Due to the increased tolerance, biofilms often pose major challenges and threats to various sectors, importantly health care (chronic infections, and antimicrobial resistance) and industry (bioremediation, agriculture, food hygiene, and antibiofouling) (1-6). On the other hand, biofilms are essential in facilitating biogeochemical processes, food digestion, immunity, biodegradation, and plant growth (1-5). Thus, given the importance and impact of biofilms, it is imperative to gain a comprehensive understanding of biofilms to limit their perils and expand on their promises.

Numerous studies have demonstrated that bacteria employ highly sophisticated physical, chemical, and biological principles to develop these stable communal forms and to establish intra- and inter-species communication across colonies (1-24). For instance, the development of biofilm invokes a complex interplay of various cellular and biochemical processes (spatiotemporal gene expression changes, gene transfer, and predation) and mechanical processes (cell-cell and cell-substrate interactions, hydrodynamic interactions, and diffusion of nutrients) acting across several length scales (7-24). The growth of bacteria in a self-imposed viscoelastic environment of EPS creates effective in-plane stresses across the entire biofilm, which, in turn, creates three-dimensional architectures (8, 9, 11-13, 15).

Despite the vast knowledge, there is still a dearth of understanding on several critical aspects of the biofilm lifestyle essential for bacterial survival and transmission. Importantly, the switching between the sessile (biofilms) to motile states, is regulated by complex and diverse mechanisms depending on the environmental signals, effectors, and signal transduction, which are yet to be unveiled (25). The action of chemical-mediated changes in motility mechanisms from motile-to-sessile or vice versa (1, 3, 4, 26) presumably altering the expression of genes that are involved in motility and biofilm formation is sparsely understood. Since motility can be the key to the success of tackling antibiotic tolerance strongly exhibited by biofilm-forming bacteria, it is imperative to identify the correlations between the chemical cues and the genetic, physiological, and molecular pathways that can activate and mediate motility.

To this end, we investigated the effects of a chemical cue, salt (NaCl, a strong inducer of stress response) on structure formation and dynamics of biofilm in an undomesticated of B. subtilis strain IITKSM1 (27) across length scales. By varying the concentration of NaCl, we observed a systematic variation in the morphology of pellicle and biofilms. While the pellicle forming ability is considerably decreased in the presence of NaCl, we observed a rapid expansion of biofilms on agar surfaces. Concomitantly, at the cellular level, NaCl induced rapid diffusive movement of bacteria along the direction of lateral expansion, in contrast to the slow super-diffusive behavior in the absence of NaCl. The rapid motility of the cells is supported by the increased flagella biosynthesis upon treatment with NaCl. Thus, our study reveals that NaCl acts as a switch that triggers the sessile-to-motile transition. Using differential gene expression analysis, we observed the upregulation of motility genes and the downregulation of genes associated with biofilm formation, thereby revealing the molecular mechanisms underlying the observed behavior. To summarize, our study demonstrates how a simple chemical cue can trigger and reprogram gene expression that modulates the dynamics of biofilms.

## Results and discussion

### NaCl-mediated regulation of pellicle formation and biofilm expansion

Figure 1 summarizes NaCl-induced changes in the formation of pellicles. A systematic increase in concentration (*W*_NaCl_) of NaCl from 0 to 2 wt.% showed drastic changes in the architecture of pellicles (Fig 1A and SF1A). We quantified these deviations by measuring the number of wrinkles (*N*_wrinkles_) and the area (*A*) covered by them (Fig 1B). Clearly, *N*_wrinkles_ and *A*/*A*_control_ (where, *A*_control_ is the area of the wrinkles in the absence of NaCl) decreased with an increase in *W*_NaCl_. The wrinkled morphology of pellicles has been observed earlier and alluded to the development of *in-plane* residual stresses due to the confined geometry over which they are growing (28). Using similar arguments, the controlled decrease in *N*_wrinkles_ with *W*_NaCl_ suggest a systematic decrease in the *in-plane* stresses developed in the pellicles. Why do the *in-plane* stresses decrease with *W*_NaCl_?

**Figure 1.**
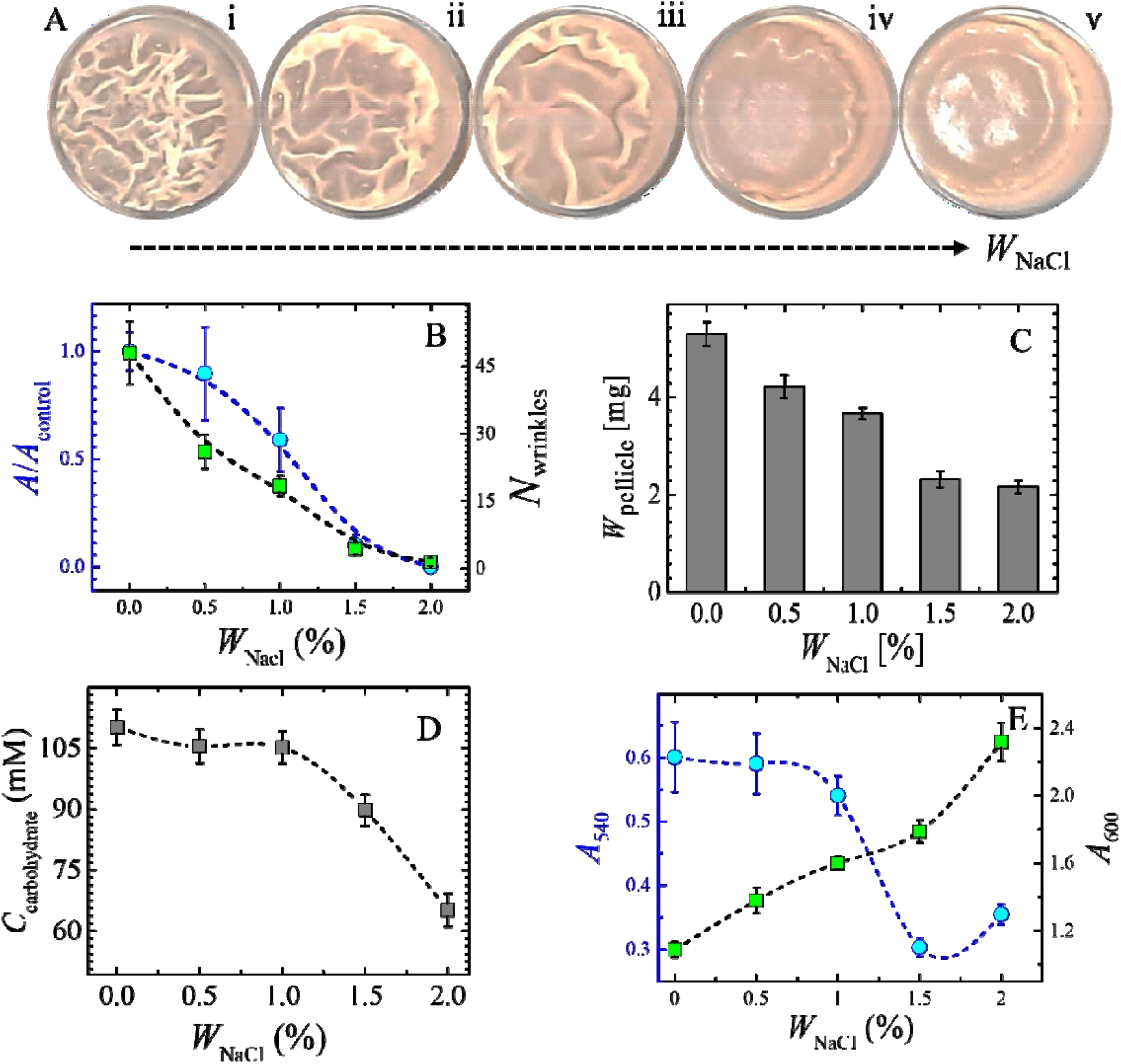
Pellicle formation in static liquid media. **A**. Top view of pellicles of *B. subtilis* IITKSM1 under different NaCl concentrations [(i)*W*_NaCl_ = 0% (ii) *W*_NaCl_ = 0.5% (iii) *W*_NaCl_ = 1% (iv) *W*_NaCl_ = 1.5% (v) *W*_NaCl_ = 2%] in liquid culture media after *ca*. 48 h of incubation at 30°C (24 well plate, Well diameter-15.5mm). All images are contrast enhanced for better visibility. **B**. Quantification of the wrinkles by plotting area covered (*A/A*control) and total number (*Nwrinkles*) as a function of *W*_NaCl_. Data plotted is the mean obtained from the biological triplicates. **C**. Dry weight of pellicles (*W*_pellicle_) as a function of *W*_NaCl_. Data plotted is the mean obtained from three independent experiments. **D**. Secreted carbohydrate concentration (*C*_carbohydrate_) was estimated using phenol-sulphuric acid method for varying NaCl concentrations. **E**. Optical density was measured for planktonic cells (*A*_600_) and crystal violet-stained surface adhered biofilm (*A*_540_) as a function of *W*_NaCl_.

To shed light on this question, we measured the overall biomass of the pellicle (*W*_pellicle_) and the growth rate of the bacteria in the presence and absence of NaCl. While there is an overall decrease in *W*_pellicle_ with increase in *W*_NaCl_ (Fig 1C and SF1B), the growth rate of bacteria (Fig SF2), and the chemical nature of the biomolecules in the pellicle remained same as shown by FT-IR spectroscopy (Fig. SF3). Concomitant to the observed decrease in *W*_pellicle_, we observed a systematic decrease in the concentration of secreted carbohydrates (*C*_carbohydrate_) with increase in *W*_NaCl_. These results are further supported by NaCl-induced reduction of the surface adhered biofilm and associated increase in planktonic growth of bacteria, as measured via the optical densities at 540 nm and 600 nm, respectively (Fig. 1E). These results suggest the possible reasons underlying the NaCl-induced deviations in the architecture of the biofilms. The overall reduction of *W*_pellicle_ may result in reduced *in-plane* stresses (28) and the decrease in *C*_carbohydrate_ suggest a possible decrease in the overall modulus of the pellicles. Assuming the buckling instability (28), the decrease in modulus of the substrate (monolayer) is expected to result in the decrease of the wavelength and, in turn, in the increased number of wrinkles. Clearly, these observations reveal that NaCl allows controlling the three-dimensional architecture of pellicles.

To improve our understanding of the importance of NaCl, we performed experiments on biofilms that are grown on agar surface and the results are summarized in Fig 2. Clearly, we observed an increase in the rate of lateral expansion of *B. subtilis* biofilms in the presence of 2 wt.% NaCl (Fig 2C), in contrast to the earlier reports showing a reduction in surface motility in the presence of NaCl (21, 29-33). As we have shown in Fig SF4, only a limited concentrations of NaCl (*W*_NaCl_ < 2.5 wt.%) showed the increased surface motility of cells, while at higher concentrations there is a decrease in surface motility consistent with a previous study (21). To further support the enhanced surface motility in the presence of NaCl, we assessed the ability of the biofilms to engulf foreign objects (34). For this purpose, we used PVDF membrane discs placed 1.5 cm apart along the lateral expansion direction. As expected, the biofilms in the presence of NaCl exhibited higher engulfing ability than the biofilms grown in the absence of NaCl (fig. 2D, 2E and fig. SF5). Evidently, the chosen low concentrations of NaCl allowed us to examine the increased motility of cells on the surface, revealing a novel characteristic of the cells in biofilms.

**Figure 2.**
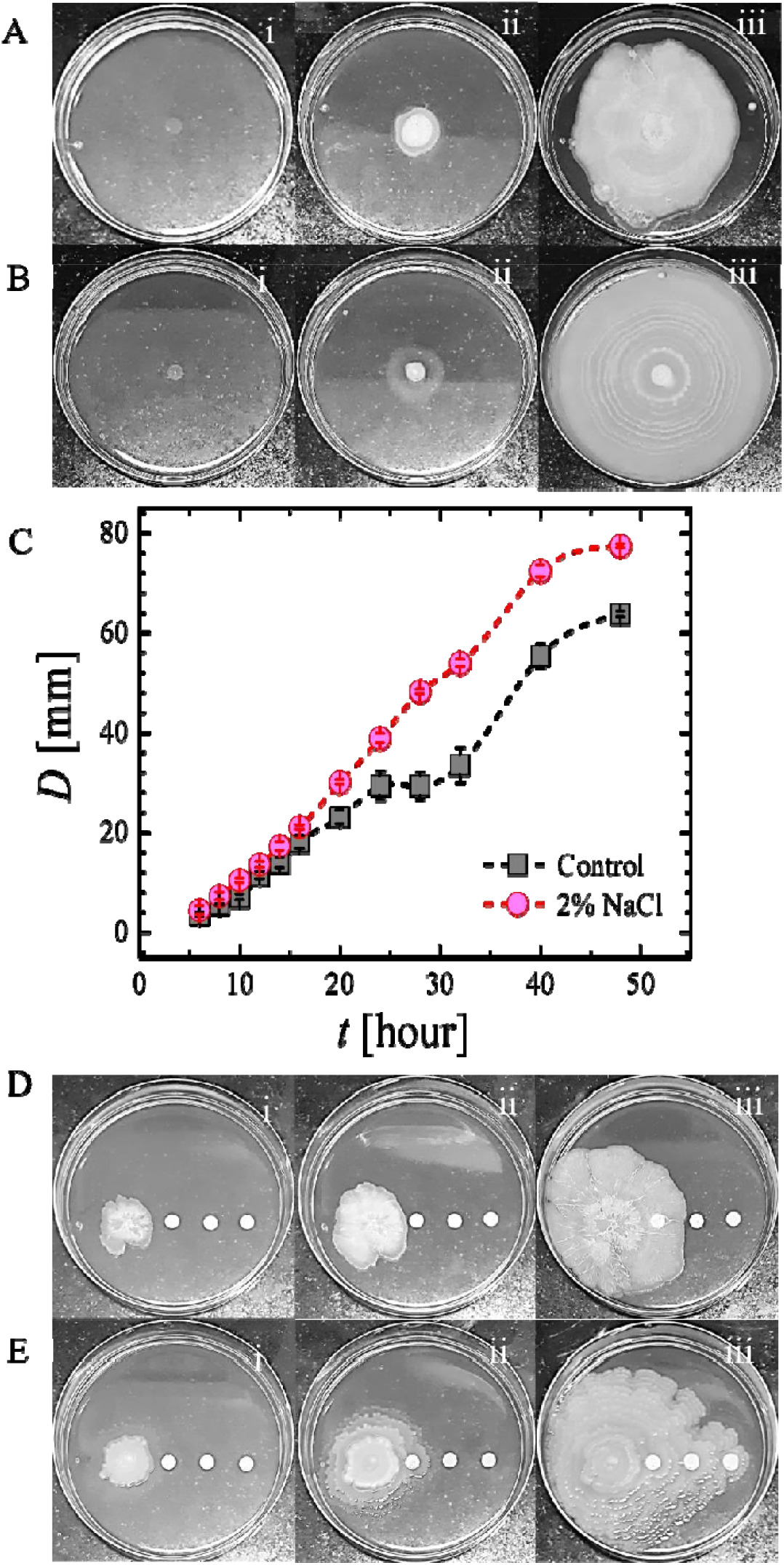
Biofilm expansion on agar surface. Representative time lapse [(i) 0 h, (ii)14 h, (iii) 24 h] images showing the lateral expansion (A) in absence of NaCl, and (B) in presence of 2 wt.% NaCl. C. Diameter (D) of the expanding biofilms (n=9) on 1.2% rich agar media Engulfment of PVDF membrane discs by B. subtilis IITKSM1 strain on 1.2% rich agar media (D) in the absence of NaCl and (E) presence of 2% NaCl. Diameter of the plates used in all the surface experiments is 90mm).

In pellicle biofilms, we observed a significant reduction in the wrinkles on adding NaCl. To verify if such variations exist in the topography of biofilms grown on the agar surface, we performed X-ray microtomography and the results are summarized in Fig 3. X-ray microtomography is a technique of choice as the matured biofilms are quite dense to be visualized by optical microscope. Assuming a chemical homogeneity at the lateral length scales of our microtomography experiments, the differences in the X-ray absorption could be related to the mean roughness of the surface. Corroborating the results of pellicle biofilms, the addition of NaCl showed a decrease in the surface roughness (Fig. 3C and 3D), suggesting the reduction in the formation of higher order structure of biofilms. Such reduction in higher-order structures is known to affect various properties of biofilms viz., penetration of foreign molecules into the biofilms (35), hydrophobicity of biofilms (36), and the mechanical tolerance of the biofilms (37-39). Here, we captured the wettability of the biofilm surface in the presence and absence of NaCl (Fig 3E and F). The contact angle of water at the center of the biofilms in the presence of NaCl was found to be *ca*. 77 °, in contrast to *ca*. 140° for biofilms formed without NaCl. Thus, biofilms grown in the presence of NaCl showed a reduced hydrophobicity, which may, in turn, be harnessed to facilitate the penetration of foreign molecules like drugs into the biofilms. These results demonstrate that NaCl controls the growth and formation of the B. *subtilis* biofilms. Does the presence of NaCl affects only at the macroscopic length scales of biofilm formation or also at the cellular and molecular scales?

**Figure 3.**
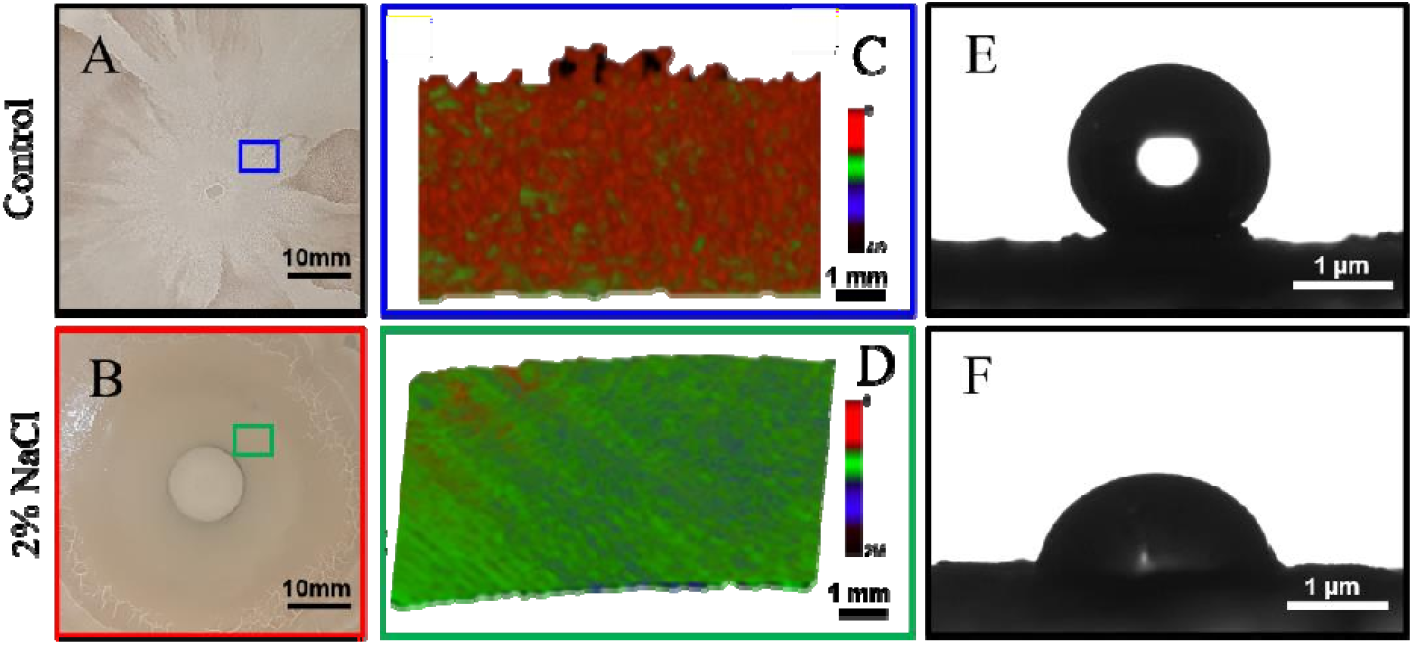
*B. subtilis* IITKSM1 colony morphology on agar with and without NaCl. Photograph focussing on the centre of a matured biofilm (**A**) in absence of NaCl and (**B**) presence of 2 wt.% NaCl. (**C**) and (**D**) X-ray Micro Tomographs corresponding the blue and green rectangles in A and B, respectively. Surface wettability of the colony centre grown (**E**) without NaCl and (**F**) with 2wt.% NaCl is captured via the contact angles of water on the respective surfaces.

### NaCl affects cell size and the kinetics of biofilm

To address this, we probed the spatiotemporal expansion of the colonies in the presence and absence of NaCl using time-lapse optical microscope. We limited our focus to the expanding edge containing a monolayer of bacteria. In Fig. 4A-C, a time series of optical micrographs were shown for biofilms, in the presence and absence of NaCl, *ca*. 8 hours after the inoculation on the agar surface. As expected from the results of Fig. 2, there is an increase in the surface motility of bacteria in the presence of 1wt.% and 2 wt.% NaCl. These results are nicely corroborated by the representative trajectories shown in the inset of Fig. 4D. To further quantify, we deduced mean squared displacements, <Δ*r*^2^(*t*)>, from the trajectories of all bacterial cells and deduced the probability distribution of maximum displacements (Figure SF7). Interestingly, we observed that the fraction of cells showing zero displacement is higher in the absence of NaCl (Figure SF7) and the collective probability (*P*_large_) for displacements, larger than at least 3.5 times the length of the bacteria, increased systematically with an increase in *W*_NaCl_ (Figure 4D). This supports an increased subpopulation of motile cells in the presence of NaCl, revealing a dynamic shift in the behavior of bacterial population of bacteria. To understand the microscopic characteristics of cellular dynamics, in Fig. 4E, we show the temporal evolution of ensemble averaged <Δ*r*^2^(*t*)> in the presence of 1 wt.% and 2 wt.% NaCl, in comparison with the control sample containing no NaCl. Overall, there is an apparent increase in the motility of cells in the presence of NaCl, corroborating our earlier results.

**Figure 4.**
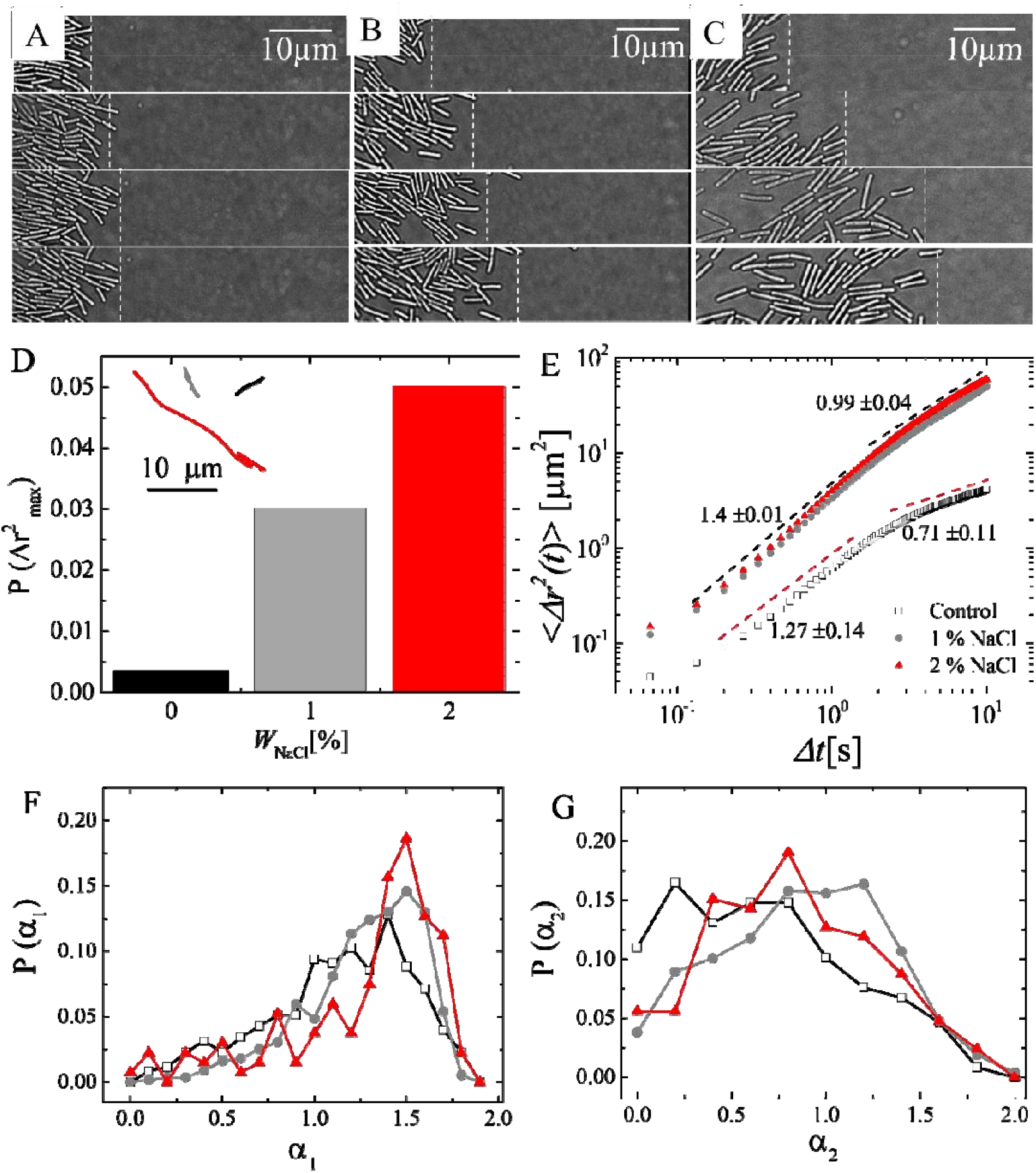
*B. subtilis* motility on agar with and without NaCl. Time series of optical micrographs after every 2-minute capturing the lateral expansion of biofilms, at the cellular level, in (**A**) the absence of NaCl, (**B**) in the presence of 1 wt.% NaCl and (**C**) 2 wt. % NaCl. Microscopy experiments were performed *ca*. 8 hours after the inoculation of culture on the agar plates. **D**. The probability that a cell has displaced at least 3.5 times larger than the length (*l*) of the bacteria i.e., (Δ ^2^| ) = ∑ (Δ ^2^| ), where 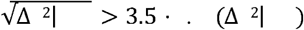 is shown as a function of fraction of _NaCl_. Inset of **D**: Representative trajectories (black and red corresponds to the trajectories in the absence and presence of NaCl, respectively) of bacteria in both the cases. **E**. Mean square displacement of motile cells in the presence of 2 wt.% NaCl (red circles), 1 wt. % NaCl (grey circles), and in the absence (black squares) of NaCl. Distribution of exponents characterizing the dynamics of bacteria, in presence and absence of NaCl, (**F**) for the initial regime (Δ*t* < 3 s) and (**G**) the later regime (Δ*t* > 3 s).

In Fig. 4E, we find two dynamic regimes, differing in the exponent α (< Δ (*t*)^2^ > ∼ *t* ^α^) characterizing the nature of microscopic dynamics. In the initial time regime, until *ca*. 3 s, the bacterial cells were displaced via super-diffusive motion in the presence and absence of NaCl, albeit with minor differences in the exponents: 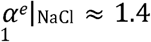 and 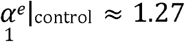. Interestingly, after *ca*. 3 s, we observed a transition from super-diffusive behavior to sub-diffusive behavior 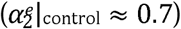 in the absence of NaCl, while the cells in the presence of NaCl displayed a transition from super-diffusive behavior to diffusive behavior 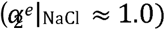. While the super-diffusive behavior of biological systems has been identified in various studies (3, 4), the observed transition from super-diffusive to diffusive (in the presence of NaCl) or sub-diffusive (in the absence of NaCl) motion is not directly apparent.

To understand these observations, we explored the values of the exponent a_1_ and a_2_, characterizing the dynamics of all the individual bacteria within the leading edge. The temporal evolution of < Δ (*t)*^2^ > corresponding to all the motile cells are shown in Fig. SF8, in comparison with the ensemble averaged ones. Interestingly, as shown in Fig. 4F and 4G, the distribution in a_1_ and a_2_ became progressively narrower as we increased the concentration of NaCl. This suggests a systematic decrease in the extent of dynamical heterogeneity with the increase in NaCl. The smaller displacements, and larger dynamic heterogeneity in the absence of NaCl indicate the existence of sub-populations of cells that are not motile, which may cause crowding of the cells. This, in turn, may result in the slowing down of dynamics as we have observed via the transition from super-diffusive motion to sub-diffusive motion. In addition, as shown in Fig. 1D, there is a larger secretion of EPS in the absence of NaCl. This may further impose limitations on the dynamics of bacteria in the absence of NaCl and support the observed transition from super-diffusive to sub-diffusive motion. On the other hand, the reduction in the concentration of secreted EPS is expected to fluidize the biofilm in the presence of NaCl. This, in addition to the reduced dynamic heterogeneity, may result in the observed transition from super-diffusive to diffusive behavior in the presence of NaCl. Moreover, it may be possible that the bacterial cells secrete other compounds in the presence of NaCl which might underlie enhanced dynamics of bacterial cells.

### Intracellular uptake of sodium and regulation of biofilm-to-motile transition

To ascertain, whether the observed phenotypic changes were due to the increase in intracellular sodium concentration, we have used inductively coupled plasma mass spectrometry (ICP-MS). The increase in the total amount of sodium inside the cells after 12, 24 and 48 hours in the presence of 2 wt.% NaCl (Fig. 5A), revealed *ca*. 6 times greater uptake of sodium. Therefore, it is highly likely that the increased intracellular levels of sodium upon NaCl treatment may be the factor governing the changes in biofilm formation of *B. subtilis*. To further examine whether the increased concentration of NaCl was responsible for the increased motility of the cells, we examined the expansion of biofilms in the presence of amiloride, a well-known inhibitor of sodium ion channel (40). As seen in Fig. 5B and 5C, 0.8 mM amiloride inhibited the motility of the cells in the presence of NaCl. Notably, both in the presence and absence of NaCl, the biofilm expansion was similar when treated with amiloride, suggesting the specific NaCl uptake resulted in the observed phenotypes. How does the cellular uptake of Na^+^ manifest into changes at the molecular level?

**Figure 5.**
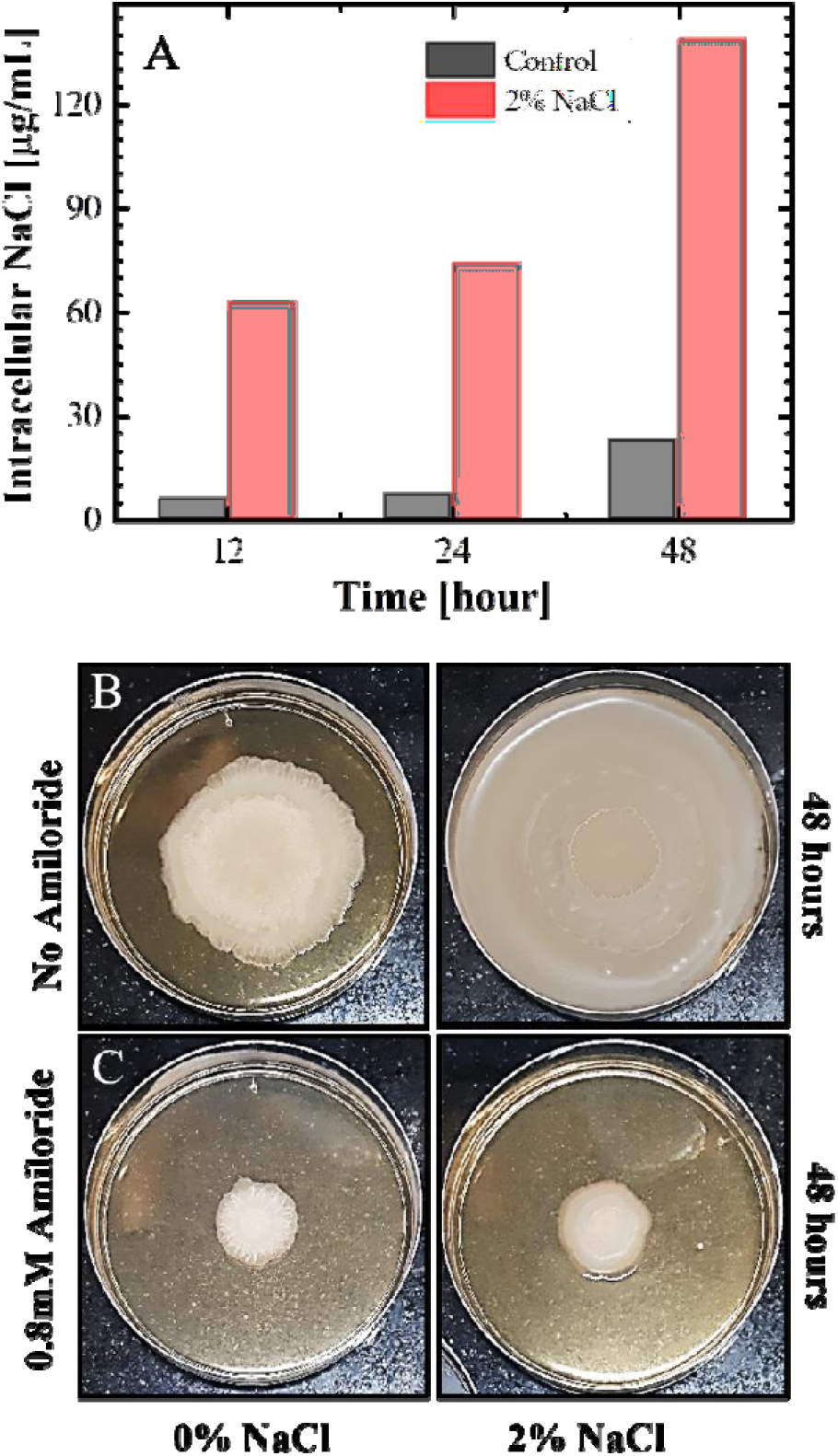
Total intracellular sodium uptake by *B. subtilis* IITKSM1 and Effect of sodium channel blocker on biofilm colony motility. **A**. Intracellular sodium concentration in the absence and presence of 2 wt.% NaCl as measured by ICP-MS. Motility of *B. subtilis* IITKSM1 on rich medium with (**B)** no NaCl and (**C)** 2 wt.% NaCl in presence and absence of 0.8 mM Amiloride, as observed after 48 hours.

To address this question, we quantified the relative change in gene expression upon the addition of NaCl, using qRT-PCR to measure the expression level of selected genes relevant for the motility of cells and the formation of biofilms. As seen in (Fig. 6A), the expression of *tapA*, encoding an important component for biofilm formation is decreased by *ca*. 4-fold. Similarly, there is *ca*. 3-fold and 2-fold decrease in the master regulator genes *spo0A* and *slrR respectively*, and *ca*. 2-fold reduction in *epsE*, the matrix producing gene. Conceivably, the reduction in the expression of *slrR, spo0A, tapA* and *epsE*, in turn, impacts (diminishes) the formation of pellicles and biofilms. On the other hand, there is *ca*. 2-fold increase in the expression of swarming gene *swrA* and chemotaxis gene *cheW* in the presence of NaCl, which likely contributes to the increased motility of the bacterial cells.

**Figure 6.**
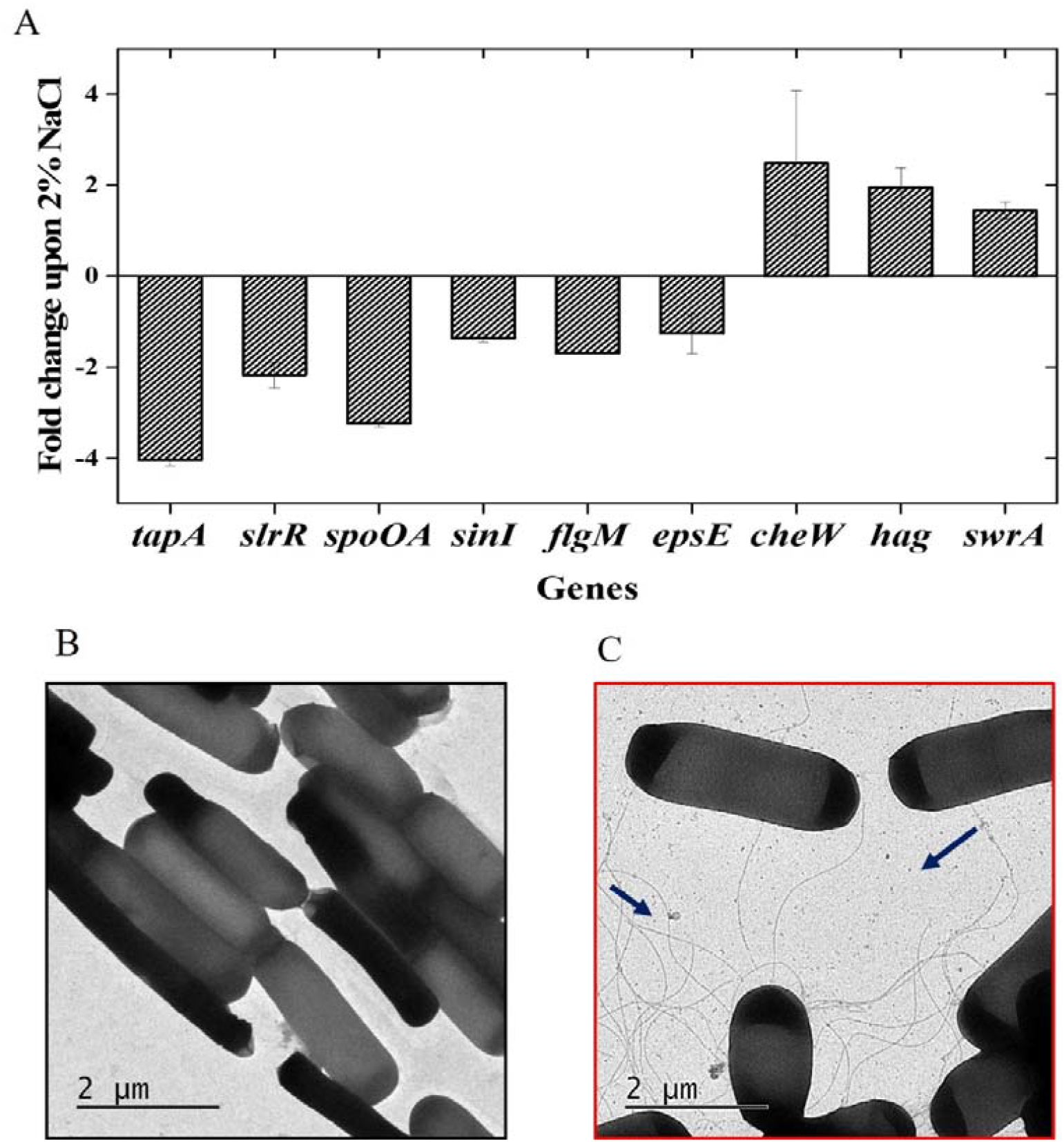
Mutually exclusive lifestyle between biofilm and planktonic state of *B. subtilis* IITKSM1 under the influence of NaCl. **A**. NaCl-induced changes in the expression level selected genes relevant for motility and biofilm formation **B**. TEM images showing cells grown without NaCl. **C**. Cells grown with 2 wt.% NaCl in liquid media. The slender structures in (C), as pointed by the dark blue arrows, show the predominant presence of flagella in the presence of 2 wt.% NaCl.

Surprisingly, in the presence of NaCl, there is *ca*. 2-fold increase in expression of *hag* gene, which is involved in the biosynthesis of flagella and there is a ca. 2-fold down-regulation of *slrR* gene that initiate the formation of biofilms by promoting cell chaining (41, 42). These changes in the gene expression, suggests the possibility of prominent differences at the level of individual cells upon NaCl treatment. To this end, we performed transmission electron microscope measurements of cells grown in the presence and absence of NaCl. As seen in fig. 6B, in the absence of NaCl, most of the cells were chained, while in the presence of NaCl (Fig 6C) the cells were less chained. These observations corroborate well with the SEM micrographs of matured biofilms, shown in Fig. SF9. Strikingly, we observed that in the presence of NaCl, most of the cells were abundantly flagellated, in contrast to the non-flagellated cells in the absence of NaCl. Thus, the enhanced motility of bacteria in the presence of 2 wt.% NaCl could be due to the reduction in chaining and induction of flagella synthesis.

### Conclusions

*B. subtilis* is a common soil-dwelling bacterium typically associated with plant roots, forming versatile biofilms with heterogeneous subpopulations. The subpopulations use a bet-hedging strategy to choose between either biofilm formation or flagella-mediated motility (43). In this study, we have characterized the importance of NaCl as a chemical cue and its potential influence the bistable switch that controls the biofilm (sessile) versus flagella-mediated motility decision. We found that a range of NaCl concentrations altered the biophysical properties, cellular morphology, gene expression and behavior of biofilms. The presence of NaCl caused a reduction in wrinkles, the dry weight and carbohydrate content of the pellicles, and reduced hydrophobicity, cumulatively indicating a major change in the biofilm architecture and properties (Figs 1-3). Salt enhanced the lateral expansion of cells and their ability to engulf foreign objects (Fig 2). These observations led to a key finding of the study, which is the increase in the percentage of a motile subpopulation within the bacterial biofilms. Cell motility was facilitated by the salt-induced increase in the flagellation of *B. subtilis* as shown by the transmission electron micrographs (Fig 6) and corroborated by the rapid diffusion of cells (Fig 4). The salt-induced changes in the expression of several relevant genes strongly supported the decision of cells to adapt motility and limit biofilm formation. Upon exposure to NaCl, the biofilm colony expresses major swarming, surfactin and flagellin genes, suggesting the possible reasons for the observed increase in the lateral expansion rates of the biofilms (Fig. 2 and 4). Simultaneously, in the presence of NaCl, we observe a decreased expression in biofilm formation genes, such as *tapA, slrR* genes and negative flagellar regulator *flgM*. Thus our data is concordant with the proposed model of a strict and reverse regulation in the transcription of biofilm matrix genes during motility (44). In addition, the significant down regulation of *epsE* in the presence of NaCl might underlie the decrease in the carbohydrate concentration (Fig. 1D) and, in turn, the changes in the architecture of the pellicle biofilms

From the standpoint of biofilm, multicellular adaptation happens via surface motility of bacteria and hence, it would be noteworthy to define how the chemical cues like NaCl act as a switch for biofilm-to-motility transition. Previous studies have discussed the impact of stressors like antibiotics and the presence of other competitive species in their ability to induce short-range or long-range orientational order of cells and bacterial motility, underlying the necessity for bacterial species to move towards more favorable niches. (1). Additionally, motility is essential to engulf foreign colonies in a competitive environment in *B. subtilis* biofilms (35). NaCl may act as a chemical cue for *B. subtilis* cells to enhance motility, which may provide competitive benefits in nature. Importantly, how chemicals typically encountered by bacterial cells in their environment influences the multicellular systems is less understood (45). Various chemicals have distinct effects on the ability of bacterial species to form biofilms. For instance, CaCl_2_ is known to inhibit surface motility and increase biofilm formation (46), while Magnesium ions are showed to inhibit biofilm formation (47). We believe that NaCl could act as a chemical trigger to enhance motility and either to promote biofilm spread or to expand to new niches. Although, biofilms are sessile, within their extracellular matrix, a subpopulation of motile cells enables biofilm spreading (3). Furthermore, salt mediated changes in the cellular states within the biofilm may act as environmental cue for dispersal of subpopulations from mature biofilms. Chemical-mediated dispersal is an active mechanism of subpopulation escaping from mature or aging biofilms. For instance, dispersal in mature *B. subtilis* is triggered by D-amino acids (48). We demonstrated that the architectural and gene expression changes induced by salt are specifically mediated by cellular uptake of salt through sodium ion channels. Overall, we present evidence that salt can reprogram gene expression, alter cellular morphology and the state of cells to adapt to motility, which may facilitate the bacterial colony escape or expansion.

Besides, our study may highlight the possibility for augmenting the efficacy of antibiotic treatments. It is well known that biofilms are refractory to antibacterial agents owing to various reasons including the sessile state and reduced diffusion of the drugs in the biofilm matrix. For instance, biofilm formation is an essential feature of *B. subtilis* that can confer resistance to antimicrobial agents (49). Our findings on NaCl-mediated biofilm-motility transition may additionally highlight an antibiotic evasion strategy. Since salt enhanced the motility of a subpopulation of cells, those motile and dividing cells can be targeted effectively by antibiotics than the sessile cells in the biofilm. To summarize, the dynamic nature of the subpopulations within bacterial biofilms and how their cellular states can be influenced purely by the chemical cues offer insights into the evolution of biofilms and their versatility in nature.

## Materials and Methods

### Bacterial strains and Media

The bacterial culture used in the study is *Bacillus subtilis* IITKSM1. The strain isolation and sequencing information are shown in our earlier work (27). The *B. subtilis* IITKSM1 was grown on rich medium containing Yeast Extract (1%), peptone (2%) and dextrose (2%). In addition to the rich medium, different NaCl concentrations ranging from 0 to 2 wt.%, amounting to a maximum molarity of *ca*. 0.37 M, were used. The pellicle formation and colony architecture assays were performed in rich media and in varied concentrations of NaCl described above. Complementary experiments were performed using minimal media glutamate glycerol (MSgg) broth as used in (50) (95.37 ml sterile MQ water, 7.5 ml 0.1 M phosphate buffer, 30 ml 0.5 M MOPS pH 7, 3 ml 0.1 M MgCl_2_, 1.05 ml 0.1 M CaCl_2_, 0.75 ml 0.01 M MnCl_2_, 0.9 ml 8.35 mM FeCl_3_, 0.15 ml 1.0 mM ZnCl_2_, 0.03 ml 0.01 M thiamine HCl, 1.5 ml 50% glycerol, 7.5 ml 10% glutamic acid, 0.75 ml 10 mg ml^−1^ tryptophan, 0.75ml 10 mgml^−1^ phenylalanine and 0.75 ml 10 mg ml^−1^ threonine 150 ml^−1^ media) or on MSgg plates supplemented with 1.5% agar and with appropriate concentrations of NaCl. All medium components were made as solutions in sterile MQ water and sterilized by either autoclaving or filter sterilization and mixed aseptically before use.). The culture was maintained on rich and LB Agar plates. The primers used in the study are listed in Table 1.

**Table 1.**
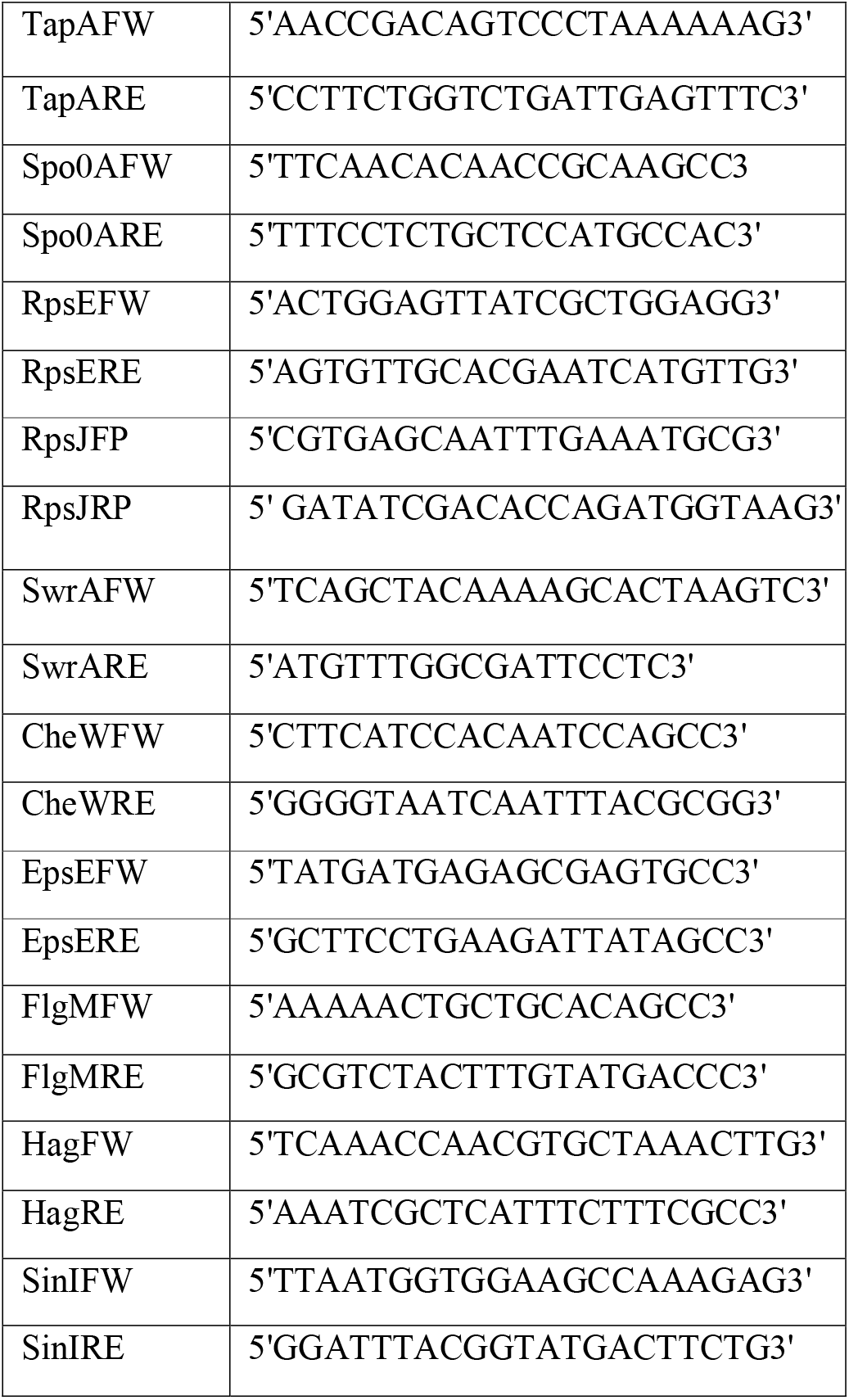

### Pellicle formation assay

For floating pellicle assays, *B. subtilis* strain IITKSM1 was cultured in rich medium and minimal medium in Tarsons (24 well) plates at 30°C for 48 hours and 72 hours, respectively. The NaCl concentrations were varied in the in each well (51). The number of pellicles, pellicle fold width and area of folds in the pellicle was analyzed using ImageJ software (52) and plotted using Origin Pro 9.1 software (OriginLab corporation [http://www.OriginLab.com]). For quantifying the dry weight of the formed pellicle, 1 ml of 100 % ethanol was carefully poured under the pellicle using a pipette to lift it from the surface of the liquid. The obtained pellicles were vacuum dried, and the weights were measured. The data shown in Fig. 1 are representative of three independent experiments.

### Surface motility Assay

Bacteria were grown in rich medium at 30 °C for 12 hours at 200 rpm. The inoculum was adjusted to obtain OD_600_∼1.0. Three microliters of cell suspension were spotted on 1.2% agar rich medium without and with 2 wt.% NaCl and grown at 30 °C for up to 48 hours. The time lapse photos were captured using straight forward mobile camera. To confirm the effect of NaCl is due to the sodium ions, a sodium channel blocker named amiloride at 0.8mM concentration in the agar medium. To observe the colony morphology in minimal MSgg media, similar procedure was followed with varying NaCl concentrations.

### Disc Engulfment assay

The extent of engulfment by *B. subtilis* IITKSM1 in presence and absence of NaCl was performed using methods discussed in reference (35). Three PVDF membrane discs of 6mm diameter were placed at 1.5 cm, 3 cm, and 4.5 cm from the inoculated biofilm colony. For each *B. subtilis* IITKSM1 colony, the number of PVDF discs engulfed by the expanding biofilm colony were captured and quantified. The results presented here are the measurements of at least 4 different colonies at each time point.

### Optical Microscopy and imaging of Motility

*B. subtilis* IITKSM1 cells swarming on surface of rich agar medium were imaged with time-lapse optical microscope (Olympus BX40 microscope) using 100X magnification (Olympus, Japan). Optical micrographs of the expanding edges, in the presence and absence of NaCl, were obtained *ca*. 14 hours after spotting on the agar plate.

### Image Analysis and Bacteria Tracking

Image analysis and cell-tracking were performed using the open access package CellProfiler [CellProfiler Project [http://www.cellprofiler.org]]. The input images were first converted to gray scale images, whose contrast is enhanced by implementing inbuilt module named “EnhanceOrSuppressFeatures” in the cellProfiler. Using the contrast enhanced image, we obtained the binary image by implementing “otsu-thresholding method”. Thresholded images as obtained are classified into cells and background using watershed algorithm. Cell orientation and its dimension were calculated by using “objectSizeShape” module in CellProfiler. Such images were subsequently used for tracking the trajectory of individual cell using a standard particle-tracking algorithm based on a “Follow neighbor criterion” in successive frames. Further, the trajectories were analyzed with python scripts to obtain mean square displacements.

### X-ray Microtomography

X-ray microtomography analysis was done using Bruker Skyscan 1275 micro-CT. A small sample (9 mm x 5 mm) was excised from the proximity of the centre of the mature biofilm on agar. The sample was placed on a rotation stage and irradiated with X-rays on the surface of the biofilm by rotating it 180° in equally spaced increments. At every angle three projections were obtained and averaged to obtain a 2D projection. The magnification chosen for the samples within the field of view which corresponded to voxel resolution of ∼ 5 μm (53).

### Analysis of Surface Wettability

The *B. subtilis* IITKSM1 cells were grown for 48 hours on rich agar medium. The contact angle measurement was determined using a goniometer (KRUSS-Drop Shape Analyzer-DSA 25E). The water droplet was placed close to the proximity of mature biofilm centre and the contact angles were measured for different samples. The drop profile was processed using the image analysis package ADVANCE software, KRUSS, GmbH, Germany.

### Transmission Electron Microscopy (TEM)

For investigation of flagella by TEM (54), *B. subtilis* IITKSM1 was grown for 12 hours in rich medium without and with 2% NaCl supplementation. The cells grown on rich medium broth for *ca*. 14 hours were absorbed onto copper grids. The grids were washed twice with PBS. Negative staining of the cells was done using 1% freshly prepared uranyl acetate. Samples were viewed in FEI Technai G2 20 twin TEM.

### RNA isolation and Q-PCR

RNA was isolated from 12-hour culture of *B. subtilis* IITKSM1grown at 37°C. The procedure was followed from (55). Briefly, 10ml of 12-hour culture was centrifuged and pelleted. The pellet was frozen and stored at −80°C for 24 hours. The pellet was later dissolved in 0.5ml of lysis buffer (30mM, 10mM EDTA with 10mg/ml lysozyme) and incubated at 37°C for 30 minutes. This was followed by addition of 1ml of Trizol reagent and 0.3ml of chloroform to the tube. The tube was centrifuged at 15000 rpm for 20 minutes at 4°C. The top aqueous layer was taken, and equal volume of isopropanol was added to it and stored at −20°C for 2 to 4 hours. The precipitated RNA was centrifuged at 15000 rpm for 20 minutes at 4°C. The obtained pellet was washed with 0.5ml with ice cold 70% ethanol. Pellets were dissolved in 20 μl double distilled water. The isolated RNA (2μg) was converted to cDNA with Quantitect Reverse Transcription kit (Qiagen, USA). The genes chosen were *epsE, tapA, spo0A, swrA, srfAA, flgM* and *cheW* with *rpsE and rpsJ* as housekeeping genes. Real time PCR was performed using Promega GoTaq Green Q-PCR Master mix, one step with the following conditions for activation of enzyme at 95°C for 30s, followed by 40 cycles of denaturation at 95°C for 30s, annealing at 55°C for 1 minute. The melt curve analysis was performed at 65°C for 1 minute in CFX connects TM Real-time PCR detection system (Bio-Rad, USA). The primers used are listed in Table 1. Relative mRNA levels were determined by fold change. This was calculated by 2- ΔΔCT using the mRNA level of rpsE and *rpsJ* which represents the house keeping gene for comparison(56).

### Quantification of planktonic cells, surface adhered Biofilm formation and amount of secreted carbohydrates

To determine how the planktonic cells and surface adhered biofilms alter under different NaCl concentrations, the *B. subtilis* IITKSM1 was grown in Tarsons (96 well) plates. To form biofilms, the cells were grown for up to 48 hours in the standardized rich medium at 30°C as static culture (57). The secreted carbohydrate concentrations were measured from the culture filtrate. The concentrations were determined with phenol sulfuric acid method using glucose as a standard (58). The data represented here are from 3 independent experiments.

### Fourier transform infrared spectroscopy (FTIR)

A mature pellicle of *B. subtilis* IITKSM1, grown for *ca*. two days on the rich medium (with and without NaCl) was cautiously separated. The pellicle was dried and lyophilized. Infrared spectroscopic measurements of the samples were performed on Bruker Tensor 27 IR spectrophotometer (Bruker Corporation, USA, KBr Beam splitter). All spectral readings were smoothened using the standard automatic smooth function (59).

### Estimation of intracellular concentration of sodium in *B. subtilis* IITKSM1 cells

Bacterial cells with balanced growth were obtained from 0% and 2% NaCl supplemented medium were pelleted, and the medium was removed. The obtained pellets were washed using 1M Tris-Cl to remove excess ions and medium without lysing the cells. The cells were lysed in 2 ml 1% HNO3 followed by sonication to break up the lysate. ICP-MS (Agilent 7900, USA) was used to estimate the intracellular concentration of potassium and sodium within the lysates. The concentrations measured were then used to estimate the average number of moles of sodium ions in a single cell, which can further be used to determine the intracellular concentration of the sodium (60).

## Supporting information

This file consists eight figures supporting the main text.

## Acknowledgments

We thank Irfan Qayoom and Ashok Kumar for allowing access to X-ray Micro CT facility, Krishnacharya Khare for contact angle measurements, Akshat Verma and Abhas singh for ICP-MS. We acknowledge the valuable suggestions from Muthukumaran Venkatachalapathy and Dharmaraja Allimuthu. We thank the funding from IITK. SM thank INSA and IYBA for the funding and fellowship, respectively.

