## Supplementary material for "NaCl Triggers the Sessile-to-Motile Transition of *Bacillus subtilis*": This file consists eight figures supporting the main text.

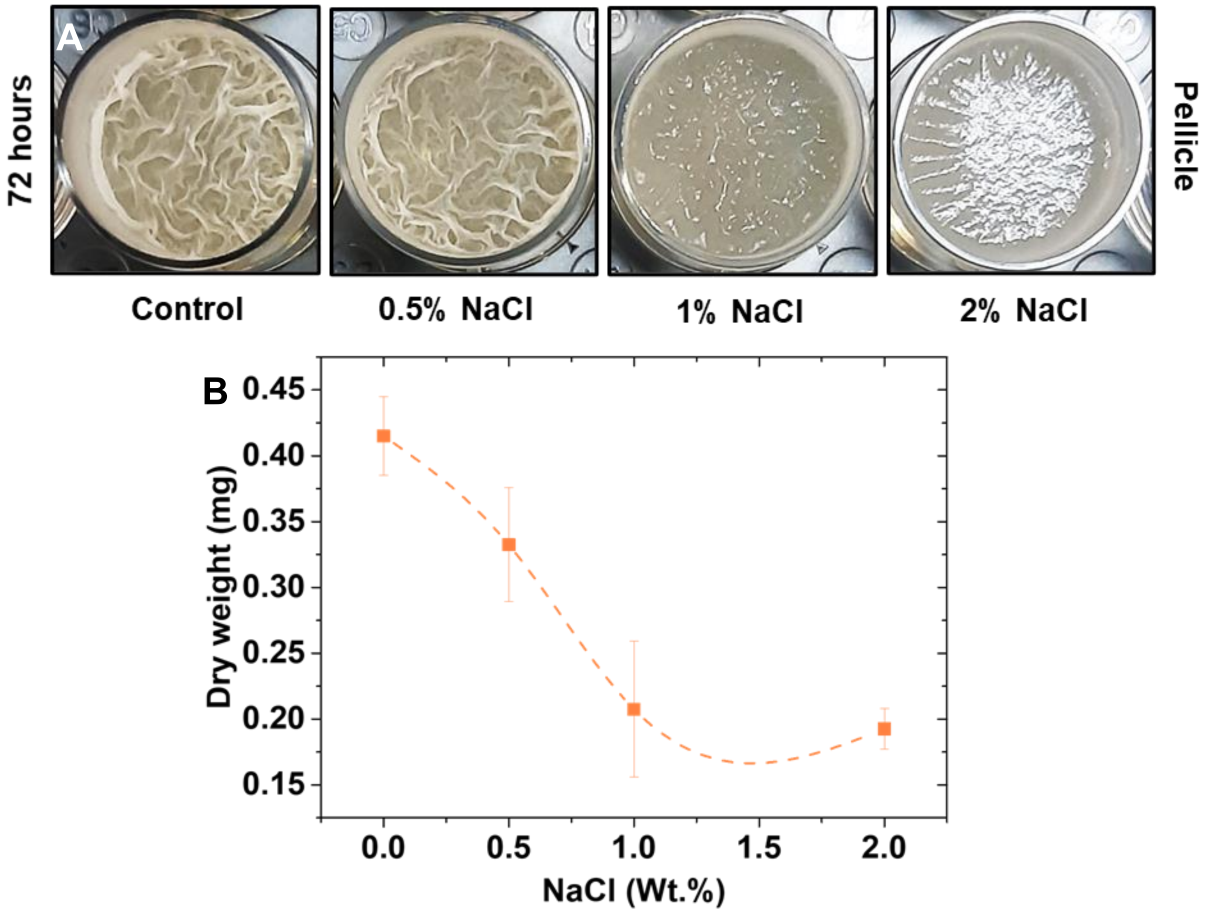


**Figure SF1. Pellicle biofilm formation in minimal media. A.** Top view of pellicle formation (24 well plate, well diameter 15.5 mm) in *B. subtilis* under different NaCl concentrations on minimal media after ~72 h of incubation at 30°C. **B.** Dry weight of pellicles were measured for each NaCl concentration from three independent experiments.

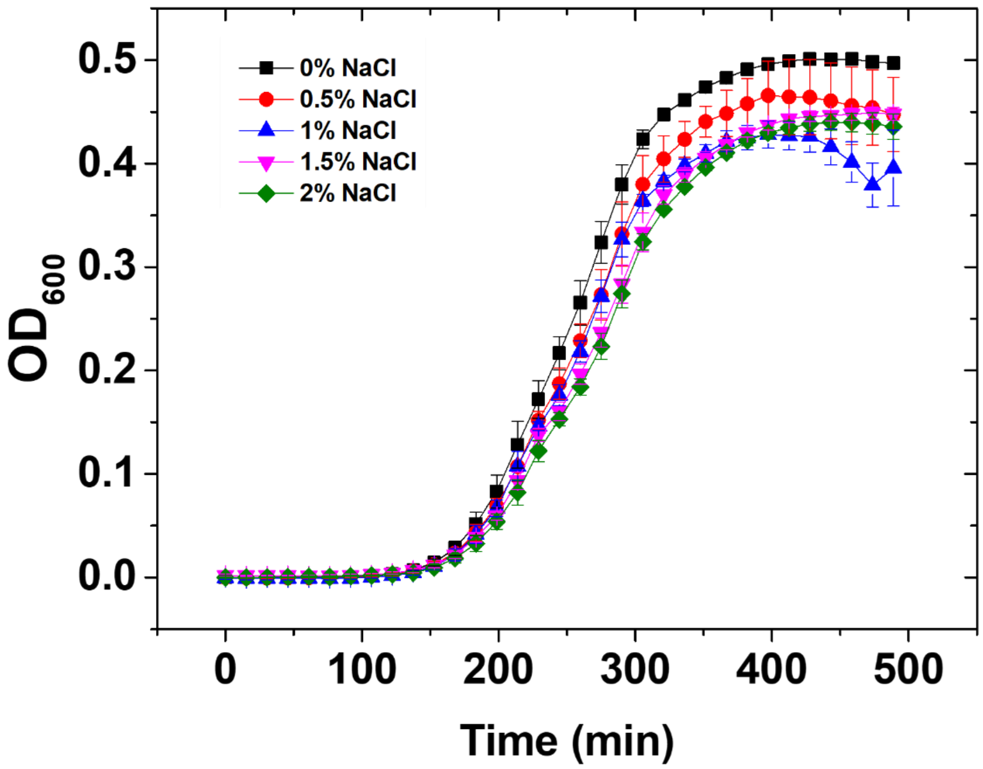


**Figure SF2. Growth of *B. subtilis* under various NaCl concentrations.** Growth curve of *B. subtilis* at 30°C with no NaCl, 0.5% NaCl, 1% NaCl, 1.5% NaCl and 2% NaCl in 96 well plate.

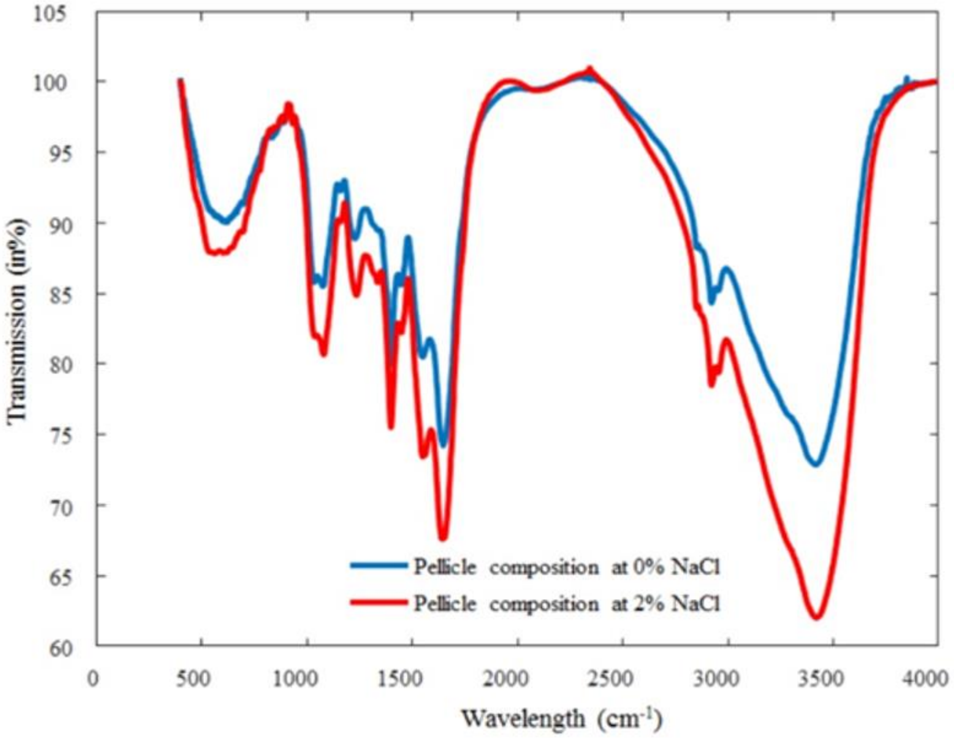


**Figure SF3. Pellicle composition of *B. subtilis.*** FTIR spectra of *B. subtilis* pellicle grown on static rich media at 30°C in the presence and absence of NaCl.

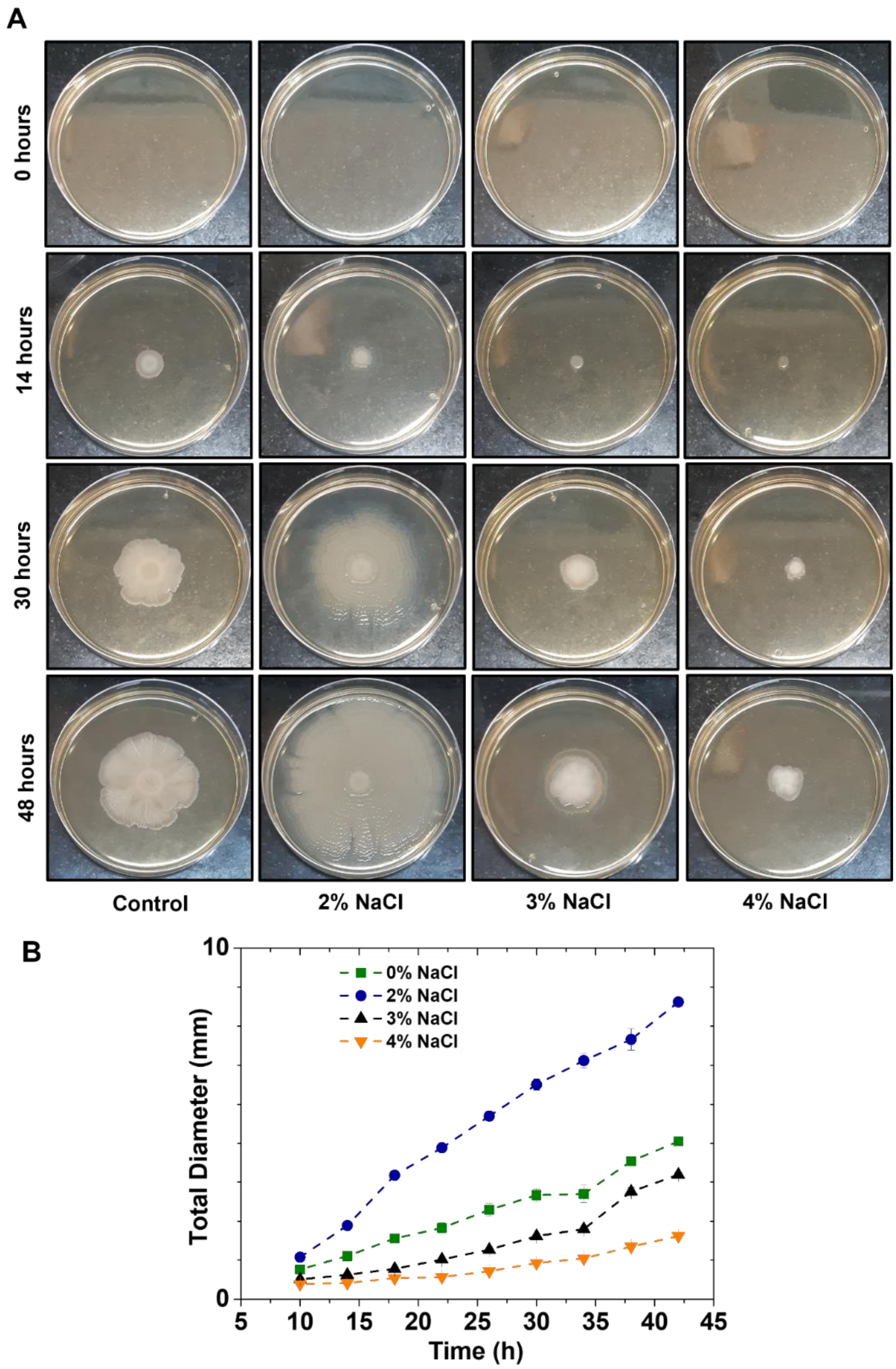


**Figure SF4**. Bacterial biofilm motility in various NaCl concentrations in 1.2% Agar. **A.** Representative images showing motility in varying NaCl concentrations. **B.** Quantification of the bacterial motility (total diameter) on 1.2% rich agar media calculated from triplicates for respective NaCl concentrations. All experiments are performed in 90 mm petri plates.

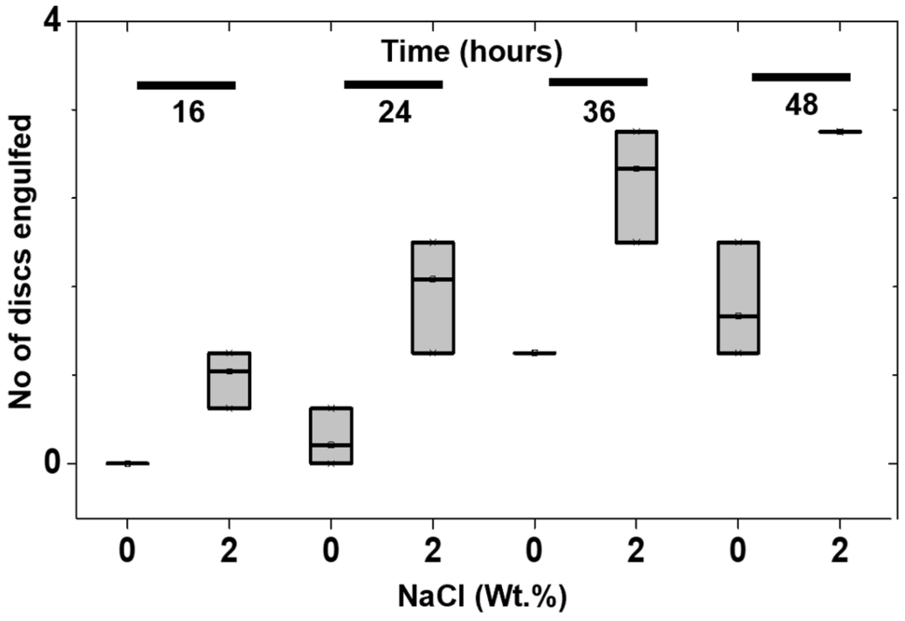


**Figure SF5.** Quantification of PVDF membrane discs engulfment ($n$ = 3) by *B. subtilis* colony in the presence and absence of NaCl treatment at various time points. The boxes indicate the upper and lower quartiles and the central line represents the median.

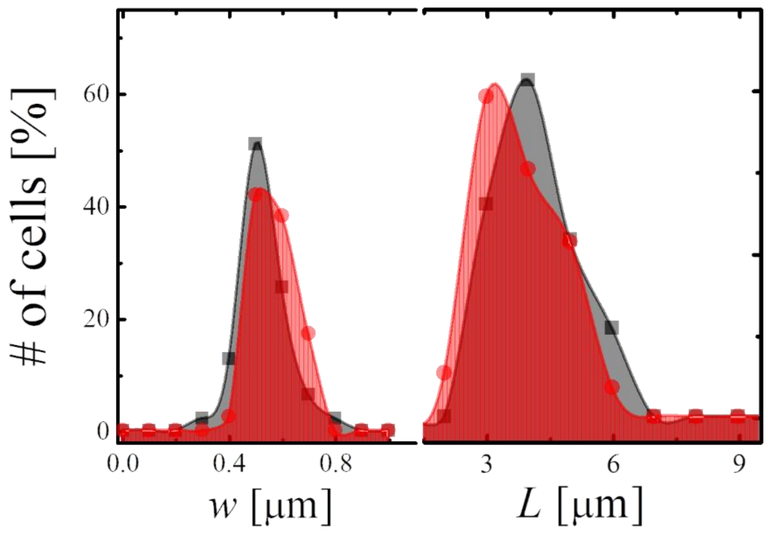


**Figure SF6**. Size of B. *subtilis* cells in the presence and absence of NaCl quantified from the images shown in Fig. 4(A) and (B) of the main manuscript. The length ($L$) and width ($w$) of cells as calculated in presence (red) and absence (grey) of NaCl.

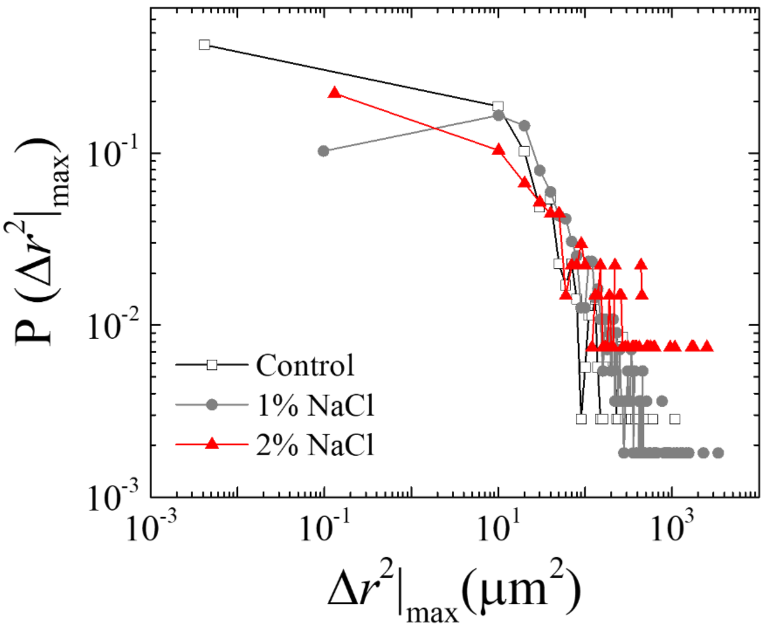


**Figure SF7.** Probability of maximum mean squared displacement of the cells in the edge of expanding colony. The data were obtained by time averaging over the images for *ca.* 30 s.

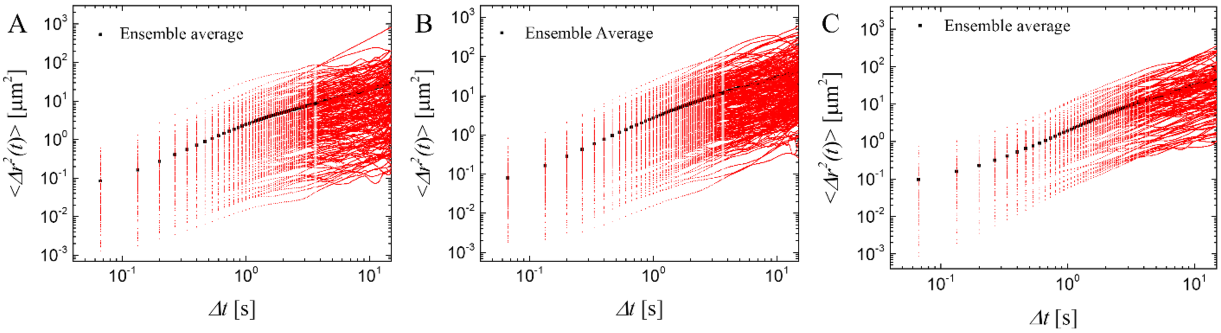


**Figure SF8.** Mean square displacement of all the motile cells (**A**) in the absence of NaCl, and in the presence of (**B**) 1wt. % NaCl and (**B**) 2wt% NaCl. Ensemble averaged data for the respective systems is shown in the corresponding figures.

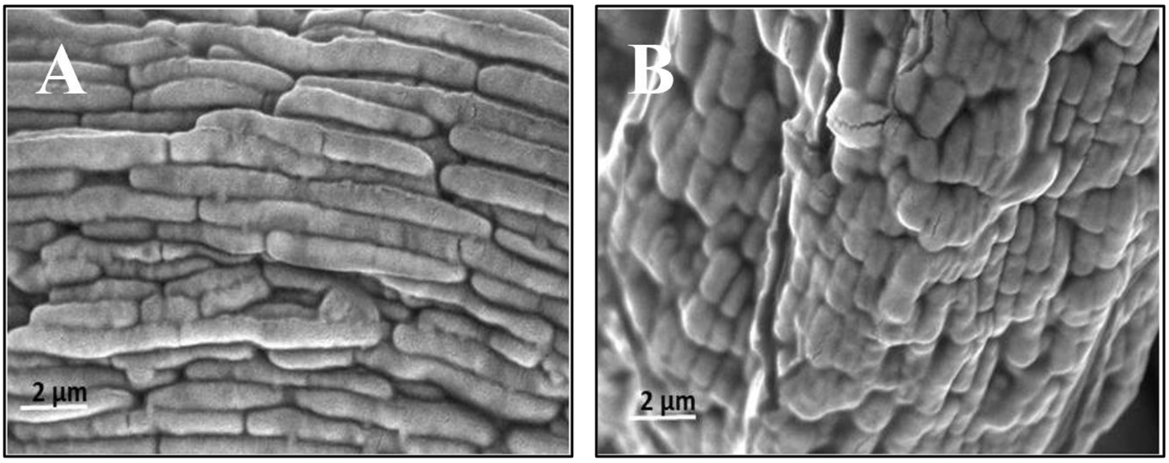


**Figure SF9**. Scanning electron micrographs capturing the cellular morphology, of a matured biofilm, in the (A) absence and (B) presence of NaCl. Cell-chaining is clearly visible in the absence of NaCl.
